# Temporal analyses of CRISPR-directed gene editing on NRF2, a clinically relevant human gene involved in chemoresistance

**DOI:** 10.1101/799676

**Authors:** Kelly Banas, Natalia Rivera-Torres, Pawel Bialk, Byung-Chun Yoo, Eric B. Kmiec

## Abstract

Recent data suggest that a time lag exists between nuclear penetration and Clustered Regularly Interspaced Short Palindromic Repeats (CRISPR)-directed gene editing activity in human cells. As CRISPR/Cas approaches clinical implementation, it is critical to establish a biological time frame in which the complex enters the cell and nucleus and executes its gene editing function. We are developing CRISPR-directed gene editing for the treatment of non-small cell lung carcinoma focusing Nuclear Factor Erythroid 2-Related Factor-Like (NRF2), a transcription factor which regulates chemoresistance. In this report, we define cellular events that surround the initialization of CRISPR-directed gene editing as a function of time. We analyze the efficiency of cellular transfection of both components of the RNP particle and assess the emergence of indels. For the first time, we image the nuclear positioning of the tracrRNA and Cas9 as a complex and as individual gene editing components. Our results indicate that while the nuclear localization of the CRISPR/Cas complex is efficient and rapid, disruption of the NRF2 gene appears four to eight hours later. We reveal an initial snapshot of the schedule and processing of CRISPR/Cas in a lung cancer cell; information that will be useful as cell-based protocols are designed and advanced.

## Introduction

Clustered Regularly Interspaced Short Palindromic Repeats (CRISPR)-directed gene editing is having a major impact on biomedical research and is already being translated into the clinic. The USA FDA has approved its use in clinical trials and similar organizations in Europe and China have done the same. The paradigm from bench to bedside has been undoubtedly accelerated because of the success and excitement that surrounds this technology. Yet, fundamental questions surrounding the mechanism of action and regulatory circuitry remain unanswered. Recently, reports have begun to appear reflecting the fact that many technological advances are not robust to be used in a broad-based format and are often reproducible only by those who discover them ^1^. To improve reproducibility and achieve technological robustness, it is important that we elucidate foundational aspects of reaction mechanics that could clearly impact the clinical application of this technology.

We have begun to investigate the feasibility of using CRISPR-directed gene editing in combination with current standard of care for the treatment of solid tumors including non-small cell lung carcinoma (NSCLC). Our initial approach has been decidedly reductionist wherein we created a CRISPR/Cas9 gene editing tool to disable the Nuclear Factor Erythroid 2-Related Factor-Like (NRF2) gene. NRF2 is a master regulator of 100-200 target genes involved in cellular responses to oxidative/electrophilic stress ^2,3^. NRF2 is also known to regulate the expression of genes involved in protein degradation and detoxification and is negatively regulated by Kelch-like ECH-associated protein 1 (KEAP1), a substrate adapter for the Cul3-dependent E3 ubiquitin ligase complex. NRF2 expression increases significantly when a NSCLC patient is treated with chemotherapy through an activation of target genes that trigger the cyto-protective response. Several studies have demonstrated that increased NRF2 expression is one of the major contributing factors of chemo-resistance in cancer cells and more prevalent in NSCLC ^3–8^. We previously discovered that successful functional knockout of the NRF2 gene in chemo-resistant lung cancer cells significantly increased the anticancer activity of cisplatin, carboplatin and vinorelbine in both cell culture and mouse models ^9^. Our results prompted testing the feasibility of innovative therapeutic approaches for clinical development to reduce clinical side effects associated with chemotherapy ^10^. This strategy centers on a combinatorial approach in which gene editing would disable the NRF2 gene removing its global regulation of chemoresistance.

Recent data suggest that a time lag exists between nuclear penetration and the appearance of gene editing activity in human cells ^11^. For clinical implementation, it is essential that a biological time frame in which the complex enters the cell, penetrates the nuclease and executes its gene editing function is established. In this report, we carry out a series of experiments to elucidate important reaction parameters that surround the coupling of nuclear penetration and gene editing activity. Our results indicate that the localization of the exogenously added CRISPR/Cas complex into the nucleus of the lung tumor cell is efficient and rapid; however, evidence of NRF2 gene disruption begins to appear after about eight hours as the gene editing components are seen exiting the nucleus. Our results provide, for the first time, a snapshot of the time-series and processing of CRISPR/Cas in a lung cancer cell, information that will be useful as clinical protocols are designed, developed and advanced.

## Results

We sought to examine the temporal relationship among nuclear localization, transit and gene editing activity vis-à-vis indel formation on the clinically relevant gene NRF2. To do so, we developed a CRISPR-directed gene editing strategy to target within the Neh2 domain of NRF2, encoded by Exon 2, using a dual fluorescently tagged CRISPR/Cas9 ribonucleoprotein (RNP) complex; the tracrRNA contained an ATTO 550 label and purified Cas9 protein was tagged with GFP (**Figures 1A and B**). After cell preparation and RNP assembly, A549 cells were transfected by Nucleofection with the dual labeled particle. After a designated period of time, we examined the targeted cell population by 1) FACS for single and dual transfection efficiency of the ribonucleoprotein components (**Figure 1C**), 2) fluorescent microscopy to visualize cellular and nuclear uptake, 3) western blot analysis for the presence and sustainability of Cas9 protein, and 4) gene editing activity on the NRF2 gene using the Tracking of Indel DEcomposition (TIDE) program ^12^.

**Figure 1.**
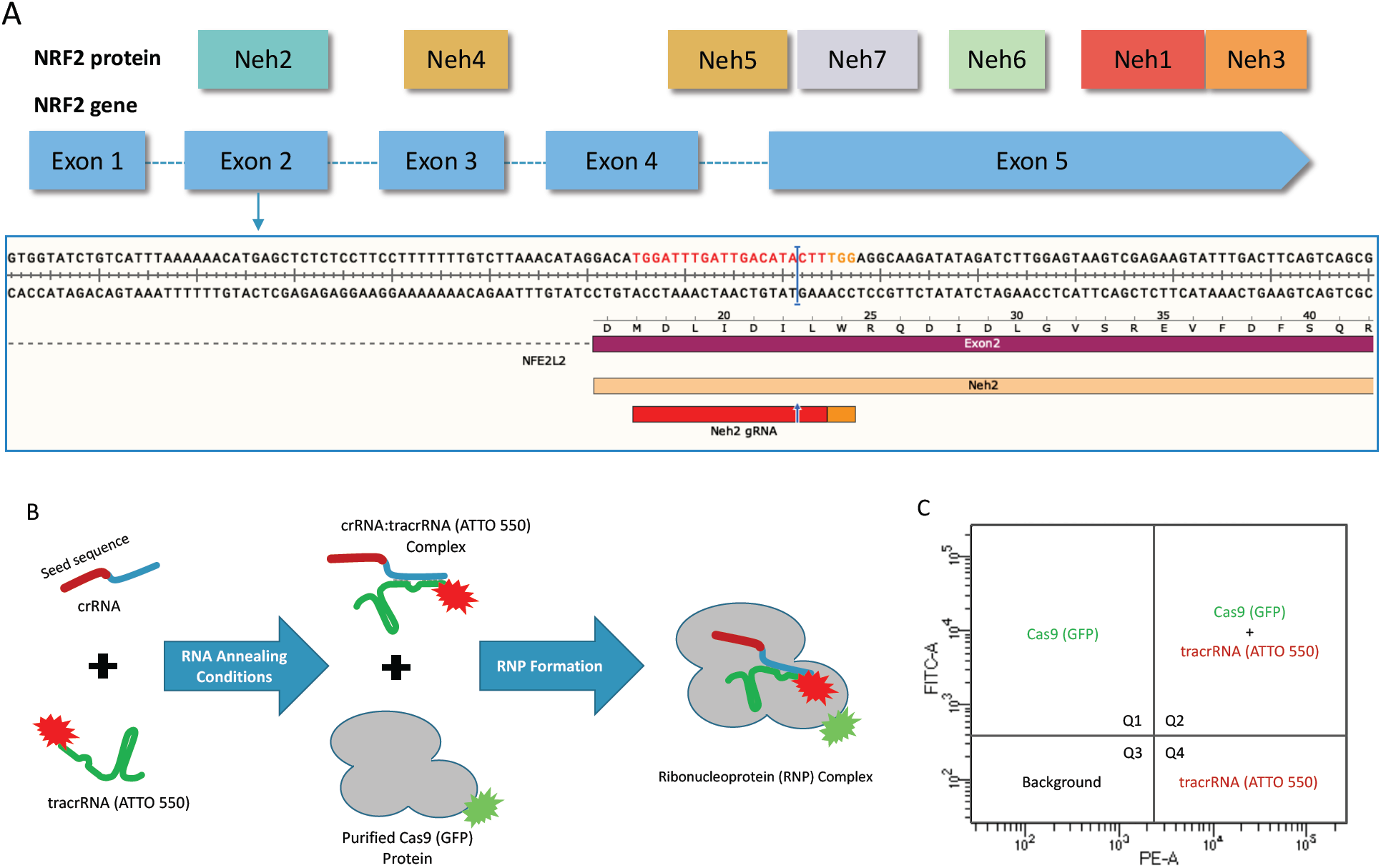
**(A)** CRISPR design and NRF2 sequence target. Structural domains of the NRF2 protein in relation to the exons of the NRF2 gene. CRISPR/Cas9 was designed to target the beginning of exon 2 using the gRNA shown in blue and the PAM sequence in red. **(B)** CRISPR/Cas9 RNP assembly reaction using fluorescently labeled components. The crRNA contains the seed sequence (shown in red) designed to target exon 2 of the NRF2 gene. The blue region of the crRNA is the interaction domain for annealing with tracrRNA (shown in green), which is labeled with the ATTO-550 dye. The crRNA and tracrRNA are annealed in equimolar concentrations. The Cas9 protein, which is labeled with GFP, is added for complete RNP formation. **(C)** FACS analysis of fluorescently labeled RNP components. An example of the localization of each fluorescent component on a FACS dot plot. Quadrant 1 (Q1) would contain cells with GFP fluorescence. Quadrant 2 (Q2) would contain cells with both GFP and ATTO 550 fluorescence. Quadrant 3 (Q3) would contain cells that do not exceed background fluorescence. Quadrant 4 (Q4) would contain cells with ATTO 550 fluorescence.

Fluorescent Activated Cell Sorting (FACS) provides an analysis of the transfection efficiency and cellular uptake by visualizing all four quadrants for co-localization of the ATTO 550 and GFP fluorescence tags respectively (**Figure 2).** The red area (P1) of each left-hand graph depicts the cells that were collected and further analyzed for fluorescent intensity (right-hand graph). The Y axis (FITC channel) represents GFP fluorescence intensity and the X axis (PE channel) represents ATTO 550 fluorescence intensity. Quadrant 1 (Q1) contains cells positive for only GFP (Cas9) while quadrant 2 (Q2), which is highlighted in green, reflects cells containing both GFP and ATTO 550 (tracrRNA) fluorescence. Quadrant 3 (Q3) contains cells that do not exceed background fluorescence and quadrant 4 (Q4) contains cells positive for only ATTO 550 (tracrRNA). The table under each plot contains raw data from each quadrant. The fluorescence (%) found in quadrant 2 is of particular interest because this population represents cells that contain both fluorescent tags, presumably a co-localization of the two components of the RNP complex.

**Figure 2.**
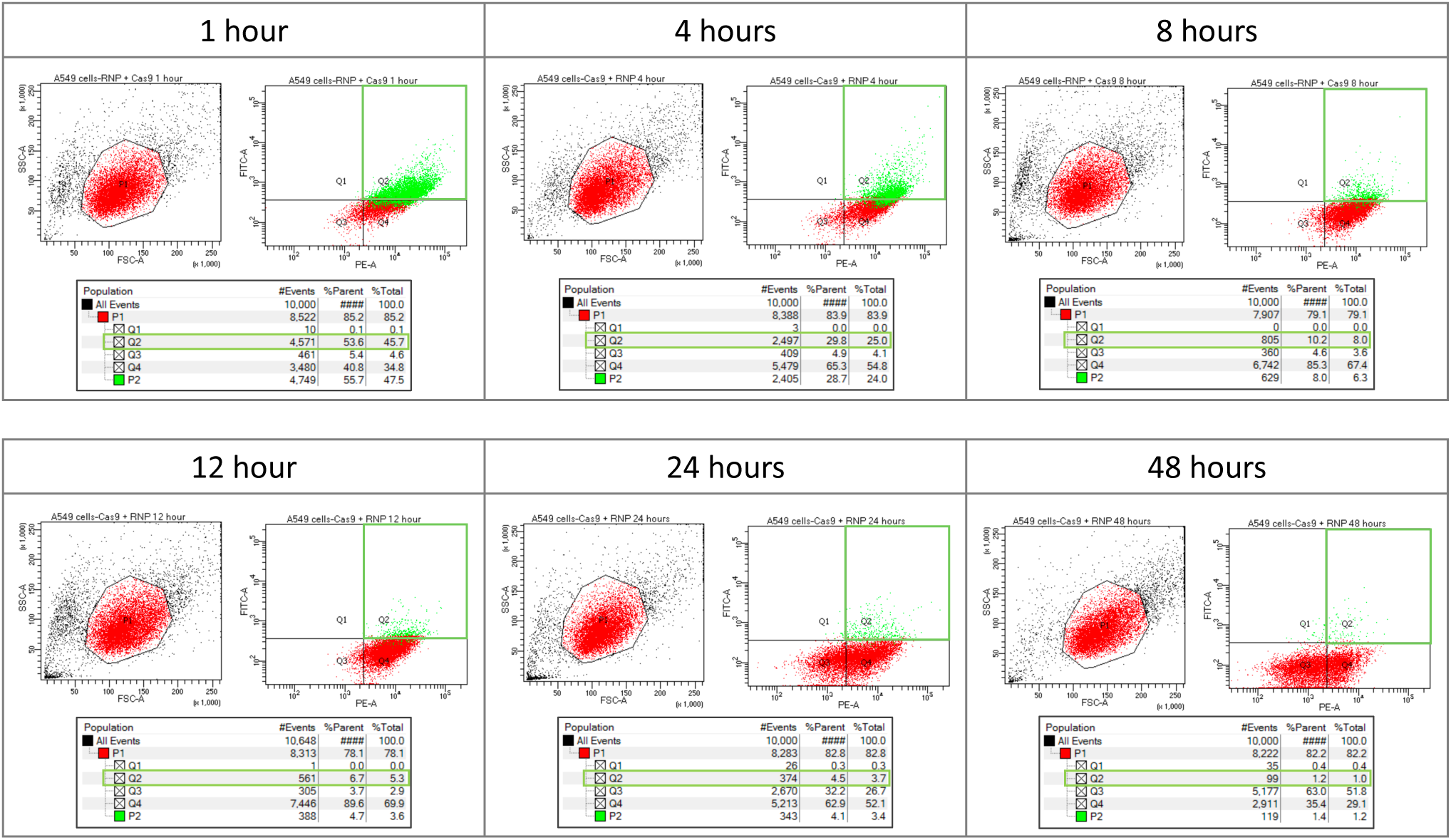
Time course analysis of fluorescently labeled RNP using FACS. A549 cells were transfected with 20 pmol of fluorescently labeled RNP complex and analyzed via FACS to determine fluorescence intensity at 1, 4, 8, 12, 24, and 48 hours post nucleofection. The red area (P1) of the left hand graph contains the population of A549 cells that were further analyzed for fluorescence intensity. Quadrant 1 (Q1) would contain cells with GFP fluorescence. Quandrant 2 (Q2) would contain cells with both GFP and ATTO 550 fluorescence. Quadrant 3 (Q3) would contain cells that do not exceed background fluorescence. Quadrant 4 (Q4) would contain cells with ATTO 550 fluorescence. A549 cells with both fluorescently labeled RNP components – Cas (GFP) and tracrRNA (ATTO550), fall within quadrant 2 (outlined in green). The table under each plot contains raw data for each quadrant.

The total number of cells with both fluorescent signals in quadrant 2 is seen to decrease over time with the maximal level observed at one-hour post nucleofection; a significant drop is seen between four and eight hours. The totality of fluorescent cells in quadrant 2 and quadrant 4 remain consistent throughout most of the reaction time (until 12 hours) indicating that the cells receiving the RNP gradually lose GFP fluorescence first, while retaining ATTO 550 fluorescence for a longer period of time. These results suggest both labels are inside the targeted cell population for approximately one to four hours. The presence of intact Cas9 protein in the cell, as viewed by western blot, confirms this time frame (**see Figure 4C**).

**Figure 3.**
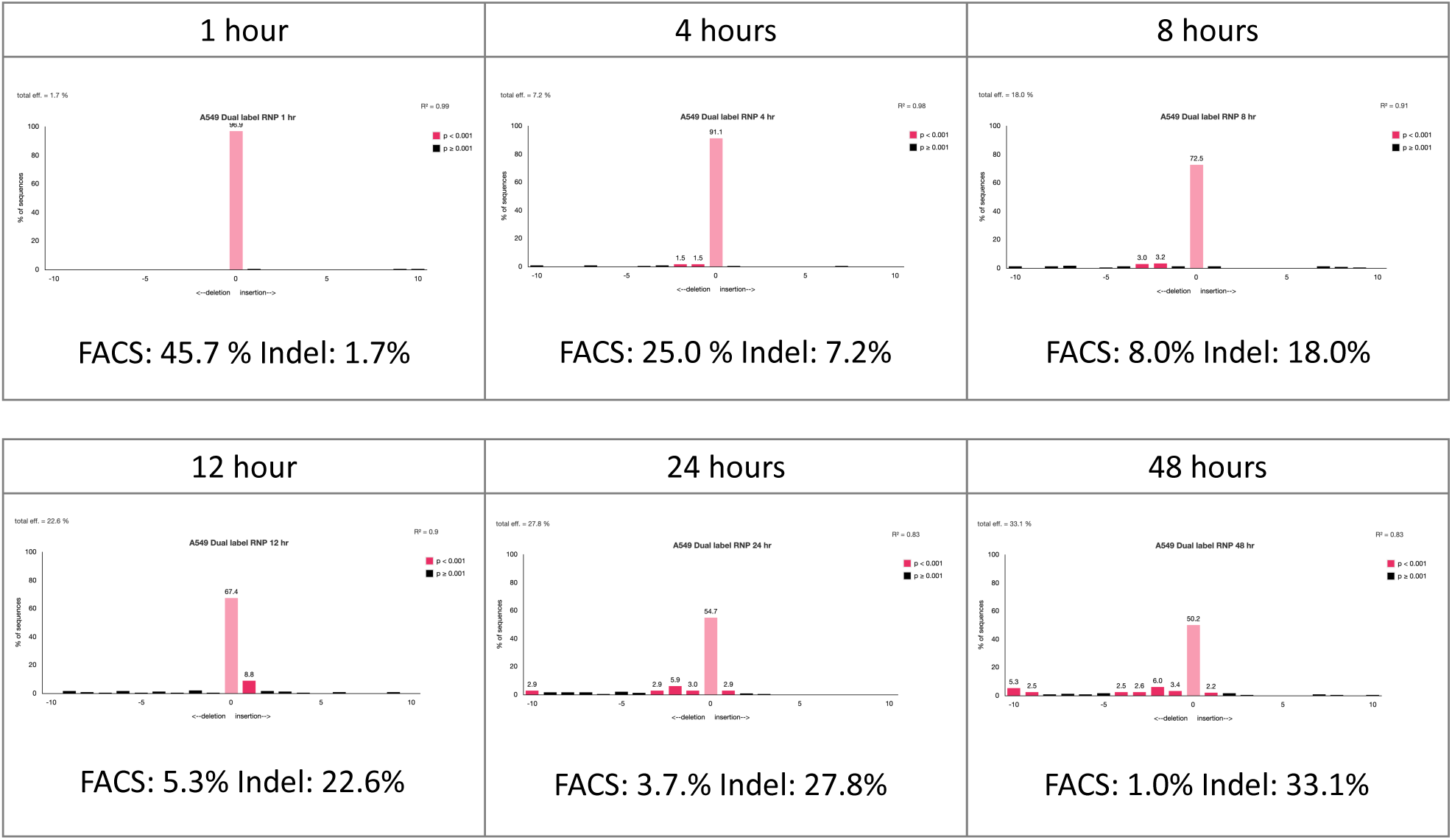
Time course analysis of gene editing activity of fluorescently labeled RNP. The total, unsorted population of A549 cells transfected with 20 pmol of fluorescently labeled RNP complex was collected as each time point. Genomic DNA was Sanger sequenced and analyzed for indel efficiency by TIDE. For comparison, the %Total of cells with both fluorescently labeled RNP components (Quadrant 2) from FACS analysis is listed adjacent to the total indel efficiency.

**Figure 4.**
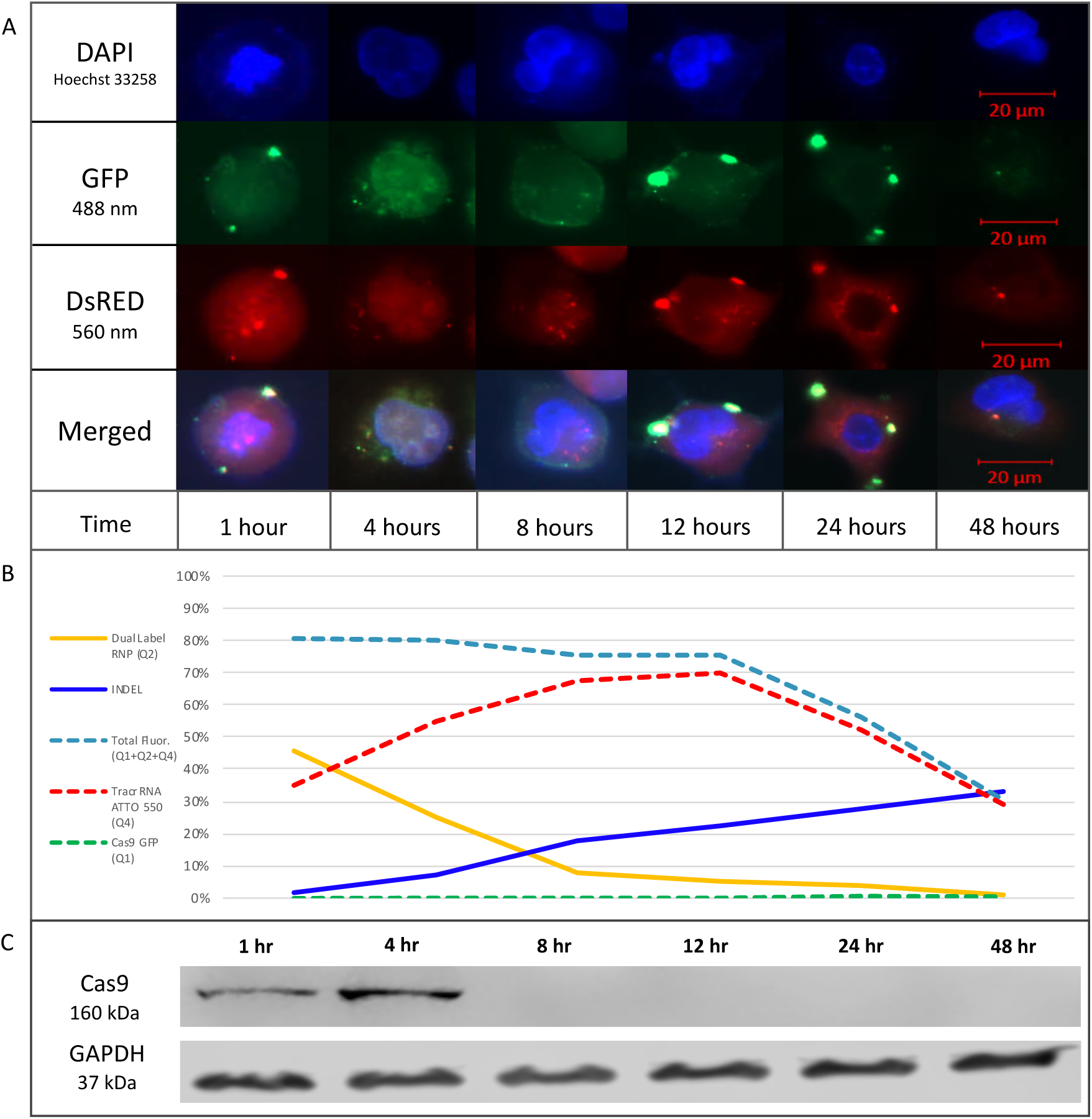
**(A)** Time course analysis of the cellular localization of fluorescently labeled RNP. A549 cells were transfected with 20 pmol of fluorescently labeled RNP complex and seeded in 4-well chambers for imaging. Representative images of each time point are shown in the figure depicting localization of the fluorescently labeled RNP complex. The brightness of the images is enhanced to better visualize the localization of the RNP complex. Scale bar represents 20 μm. **(B)** Graphical representation of individual and dual fluorescence from FACS compared to indel formation. The graph displays the %Total of each quadrant at each time point. Dual Label RNP (Q2) represents each %Total from quadrant 2, which contains cells with both fluorescent components. INDEL represents each percentage for total indel efficiency by TIDE analysis of an unsorted bulk population transfected with the dual labeled RNP. Total Fluor (Q1+Q2+Q4) represents the sum of quadrant 1, 2, and 4 to assess the total population of cells with fluorescence. TracrRNA ATTO 550 (Q4) represents each %Total from quadrant 3, which contains cells with ATTO 550 fluorescence only. Cas9 GFP (Q1) represents each %Total from quadrant 1, which contains cells with GFP fluorescence only. **(C)** Western blot analysis of Cas9 maintenance in cells. A549 cells were transfected with CRISPR/Cas9 RNP targeting Neh2 in Exon 2 and harvested at the indicated time points for western blot analysis using an antibody directed against spCas9. An antibody directed against GAPDH was used as the loading control.

To measure gene editing activity, exon 2 was amplified, Sanger sequenced and analyzed by TIDE to quantify indel formation of the total unsorted population of transfected cells **(Figure 3).** Indel formation appears to rise slowly, but steadily, throughout the course of the reaction with significant levels visible only after 8 hours, coincident with the increase in fluorescent signal of quadrant 4 of the FACS plots **(Figure 2)**.

While the timeframe of transfection efficiency and gene editing activity is informative, they are indirect measures of the RNP transit inside the cell, particularly with regard to possible nuclear localization. Therefore, we imaged nuclear entry, co-localization and exit of the RNP components by fluorescent microscopy at various time points (**Figure 4A).** Brightness and contrast are enhanced for better visualization of the fluorescent RNP components (see also **Supplemental Figure 1)**. Cells were stained with Hoechst 33258 to visualize the nucleus using the DAPI channel while fluorescent CRISPR complex components were visualized using GFP and DsRed channels respectively. The tracrRNA (ATTO 550) localizes rapidly into the nucleus while Cas9 (GFP) appears punctate at the nuclear membrane, merging or coalescing into the nucleus after four hours. Between one and eight hours, tracrRNA (ATTO 550) fluorescence appears throughout the nucleus and cytoplasm, and after eight hours, both Cas9 and tracrRNA appear to exit the cell. At twenty-four hours, Cas9 (GFP) is barely visible apart from the punctate remnants surrounding the nucleus and only small traces of tracrRNA (ATTO 550) remain visible in the nucleus. After forty-eight hours, both fluorescent components anywhere in the cell are barely detectable even with enhancement. When transfected with a single labeled RNP, GFP-fluorescing cells are located primarily in quadrant 1 after one hour and slowly disappear from all quadrants over eight hours, indicating degradation through loss of signal. Similarily, ATTO 550-fluorescing cells are observed at high levels in quadrant 4 after one hour and gradually diminish over the next twenty-four to forty-eight hours (**see Supplemental Figure 2**). Taken together, we are observing the decomposition of the CRISPR/Cas9 complex over time with Cas9 disappearing first, followed by the tracrRNA.

In **Figure 4B**, we present a graphical representation of the individual and dual fluorescence obtained from FACS analysis. Combining FACS analyses and indel formation data, the reaction could be generally broken down into two phases. Phase I encompasses RNP delivery and nuclear uptake along with the initial signals of CRISPR-directed DNA cleavage of a single target site; phase II reveals elevated levels of indel formation likely through resection and repair of the target site after cleavage, and the exit of the two fluorescent RNP components from the cell. **Figure 4C** represents Cas9 protein expression at each time point through Western blot analysis using an antibody directed against SpCas9. In our hands, the majority of Cas9 protein appears in the first four hours after introduction into the cell and becomes undetectable somewhere between four and eight hours after transfection.

## Discussion

We provide a glimpse of the cellular events that surround the initialization of CRISPR-directed gene editing of a clinically relevant gene. We analyze the efficiency of cellular transfection of an RNP particle bearing differential fluorescent labels. We also assess the emergence of indels and, for the first time, we image the nuclear positioning of the tracrRNA and Cas9 as a complex and as individual gene editing components relative to indel formation. Our results indicate that gene editing may take place in at least two definable phases. Our data suggest that Cas9 levels diminish significantly between four and eight hours which correlates to their exit from the nucleus. In contrast, the fluorescently labeled tracrRNA transits into the nucleus rapidly and remains for up to eight hours. Both fluorescent labels eventually appear as punctate particles on the nuclear membrane at twelve hours prior to being diluted and rendered undetectable between twenty-four and forty-eight hours. Quadrant 1 reveals almost no GFP fluorescence indicating that Cas9 is likely complexed with tracrRNA after introduction into the cell. Supplemental data (**Supplemental Figure 2)** confirm that the Cas9 (GFP) molecule is detectable in quadrant 1 when transfected in the absence of other fluorescent components. Fluorescence in quadrant 4 increases over twelve hours as the dual label fluorescence in quadrant 2 diminishes over that same time period. After twelve hours, total fluorescence, the sum of quadrant 1, 2 and 4 respectively, declines precipitously correlating with the images provided by fluorescent microscopy.

The initialization of genetic disruption through gene editing begins with a CRISPR/Cas complex aligning in homologous register with a target gene, facilitating site-specific double-stranded DNA breakage ^13–22^. Double stranded DNA breaks are most often repaired through a process known as Non-Homologous End Joining (NHEJ), an imperfect and unfaithful process in which nucleotides are lost or gained, generating functional knockouts. Contrasting data centered around the question of how long it takes for double-strand breaks to be created and repaired has been published with bulk measurements consistently showing that double-strand breaks have a half-life of ten to sixty minutes ^23–28^. However, these measurements lack the sensitivity that is required to follow double-strand breaks occurring and being repaired at a single locus, the reaction mechanics of CRISPR/Cas9. A crude estimate from Kim et al^29^ suggested that indel formation by CRISPR/Cas occurs early in the process but is detected only after about fifteen hours. These data align with the belief that sequence specific nucleases direct indel formation only after multiple cycles of breakage and perfect repair ^11^. Our data suggest that the time delay between penetration of the nucleus and detectable gene editing activity centers around four to eight hours, an observation that aligns with those made by Reid et al ^30^.

Multiple pathways contribute to the repair of a Cas9 induced double-strand break^31^ and relative activity of each is dependent on the locus, the metabolic state of the cell, and the half-life of the bound CRISPR complex. Indel formation likely emerges after multiple rounds of cleavage and repair once the cell has extinguished its energy supply which then causes error-prone repair of the double-strand breaks. It is possible that NHEJ initially prevents access of the Microhomology-mediated End Joining (MMEJ) pathway proteins to the double-strand break. Only after several hours wherein nonhomologous end joining has failed to repair the break, does the MMEJ pathway engage. We detect gene editing activity several hours after the RNP particle has successfully been transduced to the nucleus, a timeframe that aligns with the reaction mechanics of NHEJ and MMEJ.

Our data suggest an inverse relationship among important steps in the gene editing reaction: cellular uptake, nuclear localization of multiple components, indel formation and particle and/or component exit. These observations could help predict the timeframe of successful and impactful gene editing of the human genome. This information will be valuable to practitioners of clinical gene editing so that experimental protocols can be designed to maximize the desired outcome of genetic engineering in humans.

## MATERIALS AND METHODS

### Cell Line and Culture Conditions

Human lung carcinoma A549 cells were purchased from ATCC (#CCL-185) (Manassas, VA, USA). Cells were thawed, according to the manufacturer’s protocol, and grown in F-12K medium (ATCC, Manassas, VA, USA) supplemented with 10% fetal bovine serum (FBS) (ATCC, Manassas, VA, USA) and 1% Penicillin-Streptomycin Solution (ATCC, Manassas, VA, USA) and incubated at 37°C and 5% CO_2_.

### CRISPR/Cas9 RNP Design and Assembly

The NRF2 gene-coding sequence was entered into Benchling (https://benchling.com) and the gRNA 5’ TGGATTTGATTGACATACTT 3’ within exon 2 was selected. The GFP-labeled SpCas9 protein was a kind gift from Integrated DNA Technologies (Coralville, Iowa). Unlabeled SpCas9, ATTO 550-labeled tracrRNA and designed crRNA were purchased from Integrated DNA Technologies. RNA oligos (crRNA and tracrRNA) were mixed in equimolar concentrations to a final duplex concentration of 20μM, then annealed by heating the mix at 95 °C for 5 minutes and cooled to room temperature (15–25 °C). Prior to mixing with cells, crRNA:tracrRNA duplex (20μM working solution) and spCas9 protein (62μM stock solution) were mixed together for a final complexed concentration of 20 pmol and set to incubate at room temperature for 15 minutes.

### Transfection of Labeled CRISPR/Cas9 RNP

The Lonza SF Cell Line 4D-Nucleofector X Kit (Lonza Inc, Basel, Switzerland) was used for transfection of the RNP in the A549 cell line. For all experiments, A549 cells were seeded 48 hours prior to transfection and allowed to reach 60-80% confluency. On the day of transfection, A549 cells were harvested by trypsinization and washed twice with 1x PBS (-/-). Cells were resuspended at a concentration of 3×10^5^ cells/20μL in SF/supplement solution and 5μL of 20 pmol labeled RNP complex was added for each sample. Lonza program CM-130 was used and after 15 minutes of rest, cells were transferred to a 6-well plate or a 1.5ml tube for further analysis.

### Transfection Efficiency and Gene Editing Analysis

A549 RNP fluorescence was measured by BD FACSAria II (BD Biosciences, San Jose, CA). Cells were harvested by trypsinization, washed three times and resuspended in 1x PBS (-/-) for FACS analysis. Several controls were used to establish gates based on each fluorescent component. The controls consisted of mock transfection of A549 cells, transfection of RNP with Cas9 (GFP), and transfection of RNP with tracrRNA (ATTO 550). Gates were established based on these parameters. After FACS analysis, cellular genomic DNA was isolated from each sample using the DNeasy Blood and Tissue Kit (Qiagen, Cat. 69506). The region surrounding the CRISPR target site was PCR amplified using Q5 High-Fidelity 2X Master Mix (New England BioLabs, Cat. M0492) (762 bp, forward primer 5’-ATTAAACAAGGGTGGGATTTCTTCTC-3’, reverse primer 5’-AACTCAGGTTAGGTACTGAACTCATCA-3’). The PCR reaction was purified using the QIAquick PCR Purification Kit (Qiagen, Cat. 28106) and sent out to GENEWIZ, LLC (South Plainfield, NJ) for Sanger Sequencing. The software program, TIDE, was used for bulk sequence analysis to assess indel efficiency ^12^.

### Western Blot Analysis

A549 cells were transfected with unlabeled Cas9 RNP as described above. Cells were harvested at the indicated time points and the western blot was carried out as previously described (Bialk et al). Primary antibody incubation was performed overnight on a shaker at 4°C for Cas9 (1:5,000, Abcam ab210752), and GAPDH (1:10,000, Abcam ab8245), and secondary antibody (Jackson ImmunoResearch Laboratories, West Grove, PA, USA) incubations were all done 1 hr at room temperature at a 1:10,000 dilution. The protein bands were visualized via chemiluminescence using a SuperSignal West Dura Extended Duration ECL (Pierce) and detected on the LI-COR Odyssey FC.

### Fluorescence Microscopy

Transfected A549 cells were seeded in 4-well chambers (LabTek II). Before imaging at each indicated time point, cells were washed twice with PBS (-/-) and incubated with Hoechst 33258 (1:10,000, Invitrogen, Cat. H3569) for a minimum of 15 minutes at room temperature in the dark. Chambers were imaged at 20x magnification using the DAPI (358 nm), GFP (488 nm), and DsRed (560 nm) channels on the Zeiss Axio fluorescent observer.Z1 microscope. Random fields were imaged and images were processed on the AxioVision software. The brightness and contrast of the images have been enhanced for better visualization.

## Acknowledgments

We thank the members of the Kmiec laboratory for input and advice. We thank Dr. Lynn Opdenaker at the CTCR Flow Cytometry Core Facility for running, analyzing, and sorting cells. This project was supported by the Delaware INBRE and COBRE programs, with grants from the National Institute of General Medical Sciences – P20 GM103446 and P20 GM109021 from the NIH and the State of Delaware. This content is solely the responsibility of the authors and does not necessarily represent the official views of the NIH.

## References

1. Garde, D. A richly funded CRISPR alternative has a reproducibility problem. STAT (2019). Available at: https://www.statnews.com/2019/07/26/a-richly-funded-crispr-alternative-has-a-reproducibility-problem/. (Accessed: 23rd September 2019)

2. Chen, J., Solomides, C., Simpkins, F. & Simpkins, H. The role of Nrf2 and ATF2 in resistance to platinum-based chemotherapy. Cancer Chemother. Pharmacol. 79, 369–380 (2017).

3. Wang, X.-J. et al. Nrf2 enhances resistance of cancer cells to chemotherapeutic drugs, the dark side of Nrf2. Carcinogenesis 29, 1235–43 (2008).

4. Frank, R. et al. Clinical and Pathological Characteristics of KEAP1-and NFE2L2-Mutated Non-Small Cell Lung Carcinoma (NSCLC). Clin. Cancer Res. 24, 3087–3096 (2018).

5. Singh, A. et al. Dysfunctional KEAP1-NRF2 interaction in non-small-cell lung cancer. PLoS Med. 3, e420 (2006).

6. Hayden, A. et al. The Nrf2 transcription factor contributes to resistance to cisplatin in bladder cancer. Urol. Oncol. Semin. Orig. Investig. 32, 806–814 (2014).

7. Tian, Y. et al. Modification of platinum sensitivity by KEAP1/NRF2 signals in non-small cell lung cancer. J. Hematol. Oncol. 9, 83 (2016).

8. Torrente, L. et al. Crosstalk between NRF2 and HIPK2 shapes cytoprotective responses. Oncogene 36, 6204–6212 (2017).

9. Bialk, P., Wang, Y., Banas, K. & Kmiec, E. B. Functional Gene Knockout of NRF2 Increases Chemosensitivity of Human Lung Cancer A549 Cells in Vitro and in a Xenograft Mouse Model. Mol. Ther. - Oncolytics 11, 75–89 (2018).

10. Hirsch, F. R., Suda, K., Wiens, J. & Bunn, P. A. New and emerging targeted treatments in advanced non-small-cell lung cancer. Lancet 388, 1012–1024 (2016).

11. Brinkman, E. K. et al. Kinetics and Fidelity of the Repair of Cas9-Induced Double-Strand DNA Breaks. Mol. Cell 70, 801-813.e6 (2018).

12. Brinkman, E. K., Chen, T., Amendola, M. & van Steensel, B. Easy quantitative assessment of genome editing by sequence trace decomposition. Nucleic Acids Res. 42, e168.-(2014).

13. Barrangou, R. & Doudna, J. A. Applications of CRISPR technologies in research and beyond. Nat. Biotechnol. 34, 933–941 (2016).

14. Prakash, V., Moore, M. & Yáñez-Muñoz, R. J. Current Progress in Therapeutic Gene Editing for Monogenic Diseases. Mol. Ther. 24, 465 (2016).

15. Carroll, D. Genome editing: progress and challenges for medical applications. Genome Med. 8, 120 (2016).

16. Doudna, J. A. Genomic Engineering and the Future of Medicine. JAMA 313, 791 (2015).

17. Doudna, J. A. & Charpentier, E. The new frontier of genome engineering with CRISPR-Cas9. Science (80-.). 346, 1258096–1258096 (2014).

18. Hsu, P. D., Lander, E. S. & Zhang, F. Development and applications of CRISPR-Cas9 for genome engineering. Cell 157, 1262–78 (2014).

19. Komor, A. C., Badran, A. H. & Liu, D. R. CRISPR-Based Technologies for the Manipulation of Eukaryotic Genomes. Cell 168, 20–36 (2017).

20. Staahl, B. T. et al. Efficient genome editing in the mouse brain by local delivery of engineered Cas9 ribonucleoprotein complexes. Nat. Biotechnol. 35, 431–434 (2017).

21. Zetsche, B. et al. Cpf1 Is a Single RNA-Guided Endonuclease of a Class 2 CRISPR-Cas System. Cell 163, 759–771 (2015).

22. Jinek, M. et al. A programmable dual-RNA-guided DNA endonuclease in adaptive bacterial immunity. Science 337, 816–821 (2012).

23. DiBiase, S. J. et al. DNA-dependent protein kinase stimulates an independently active, nonhomologous, end-joining apparatus. Cancer Res. 60, 1245–53 (2000).

24. Metzger, L. & Iliakis, G. Kinetics of DNA Double-strand Break Repair Throughout the Cell Cycle as Assayed by Pulsed Field Gel Electrophoresis in CHO Cells. Int. J. Radiat. Biol. 59, 1325–1339 (1991).

25. Núñez, M. et al. Radiation-induced DNA double-strand break rejoining in human tumour cells. Br. J. Cancer 71, 311–316 (1995).

26. Schwartz, J. L., Rotmensch, J., Giovanazzi, S., Cohen, M. B. & Weichselbaum, R. R. Faster repair of DNA double-strand breaks in radioresistant human tumor cells. Int. J. Radiat. Oncol. 15, 907–912 (1988).

27. Stenerlöw, B., Karlsson, K. H., Cooper, B. & Rydberg, B. Measurement of prompt DNA double-strand breaks in mammalian cells without including heat-labile sites: results for cells deficient in nonhomologous end joining. Radiat. Res. 159, 502–10 (2003).

28. Wang, M. et al. PARP-1 and Ku compete for repair of DNA double strand breaks by distinct NHEJ pathways. Nucleic Acids Res. 34, 6170–6182 (2006).

29. Kim, S., Kim, D., Cho, S. W., Kim, J. & Kim, J.-S. Highly efficient RNA-guided genome editing in human cells via delivery of purified Cas9 ribonucleoproteins. Genome Res. 24, 1012–1019 (2014).

30. Reid, D. A. et al. Organization and dynamics of the nonhomologous end-joining machinery during DNA double-strand break repair. Proc. Natl. Acad. Sci. 112, E2575–E2584 (2015).

31. Bothmer, A. et al. Characterization of the interplay between DNA repair and CRISPR/Cas9-induced DNA lesions at an endogenous locus. (2017). doi:10.1038/ncomms13905

