## Supplemental Figures for "Temporal analyses of CRISPR-directed gene editing on NRF2, a clinically relevant human gene involved in chemoresistance"

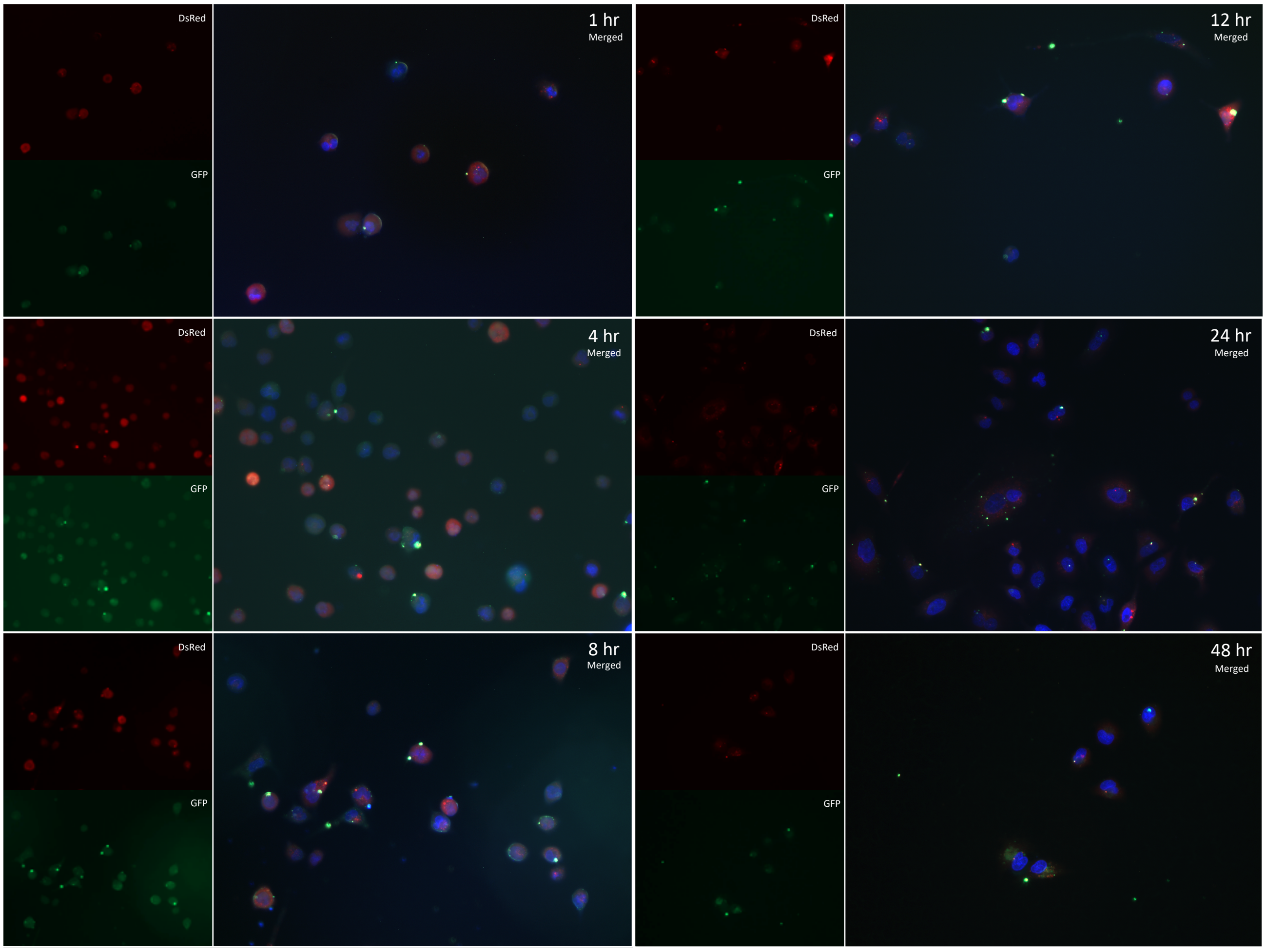


**Figure S1.** Representative images used for time course analysis. The series of images represents the average field seen for each time point. Brightness has been enhanced for better visualization of fluorescence.

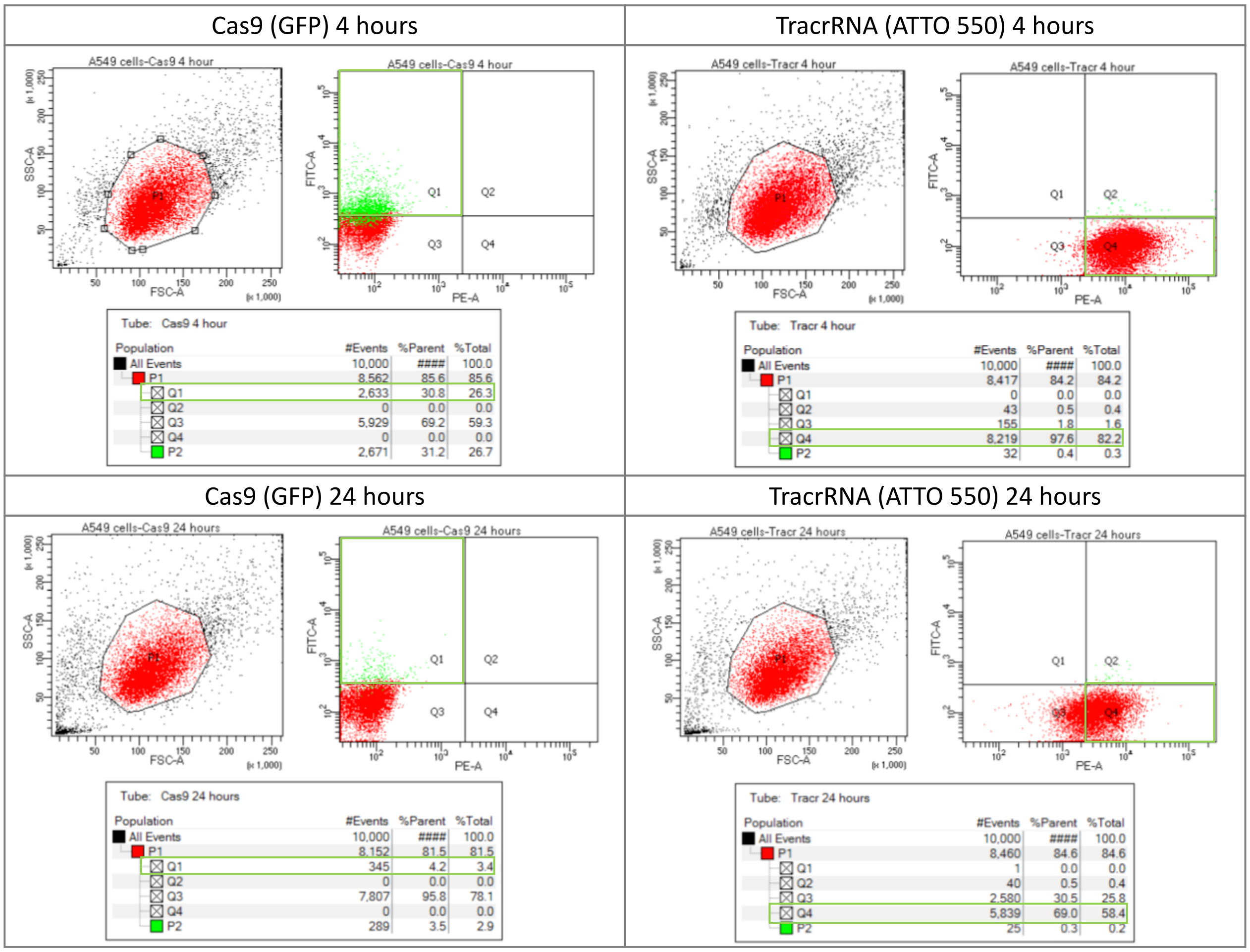


**Figure S2.** FACS Analysis of A549 cells transfected with CRISPR/Cas9 RNP containing a single fluorescent component. Representative FACS plots depicting the population of cells transfected with an RNP containing either fluorescent component at the indicated time points (data for 1, 8, 12, 48 hours not shown). Each single fluorescent component was used to set gates for dual and background fluorescence as seen in Figure 2.
